# Cell-type-specific adaptations to mitochondrial stress underly the neurological presentations of *MTRFR* mutations

**DOI:** 10.64898/2025.12.30.696998

**Authors:** Mariana Zarate-Mendez, Kaela O’Connor, Natalia Malig, Oliver Podmanicky, Julia Kleniuk, Sally Spendiff, Evan Reid, Hanns Lochmuller, Denisa Hathazi, Rita Horvath

**Author notes:** Corresponding authors: Rita Horvath, MD, PhD, Denisa Hathazi, PhD, Department of Clinical Neurosciences, University of Cambridge, John Van Geest Centre for Brain Repair, Robinson Way, CB2 0PY, Cambridge, UK. Joint last authorship (Denisa Hathazi, Rita Horvath).

## Abstract

Mitochondrial diseases are a group of heterogeneous genetic disorders that exhibit striking tissue specificity. Neurological involvement is among the most consistent features, yet the mechanisms that determine why selective neuronal populations are particularly vulnerable to mitochondrial dysfunction remain poorly understood. Mutations in *MTRFR*, a mitochondrial ribosome rescue factor, cause a progressive neuromuscular phenotype, but no relevant disease model exists to explain its cell-type-specific pathology. Here, we established the first human iPSC-derived neuronal model of MTRFR loss and identified mechanisms driving differential vulnerability between cortical and motor neurons. Although knockdown led to comparable deficits in mitochondrial translation and OXPHOS across both subtypes, cortical neurons engaged adaptive programs, including dendritic mitochondrial remodelling and heat-shock response activation, that preserved survival. Motor neurons failed to mount these responses and instead displayed apoptotic and inflammatory priming. Pharmacological enhancement of stress adaptation rescued motor neuron survival, indicating that resilience is programmable. These findings provide the first mechanistic evidence that neuronal susceptibility to mitochondrial translation defects is defined by the capacity to activate mitochondrial and cytoprotective stress-response pathways.

## INTRODUCTION

Mitochondria are central to neuronal function, integrating energy metabolism with signalling and survival pathways. Because neurons are highly polarised and have limited energy reserves, they rely almost entirely on local mitochondrial ATP synthesis to sustain activity (Rangaraju et al. 2014). Their distinctive architecture and high metabolic demand make them particularly vulnerable to mitochondrial dysfunction. Mitochondrial mechanisms contribute to neurodegenerative conditions, including Parkinson’s disease and amyotrophic lateral sclerosis (ALS) (Wood et al. 2021; Monzio Compagnoni et al. 2020; Stykel et al. 2025), where only certain neuronal populations are affected. The mechanisms that determine why some neuronal subtypes are more susceptible to mitochondrial stress while others remain resilient are not well understood. Defining the molecular basis of this selective vulnerability is essential for identifying strategies to enhance neuronal resilience.

Mitochondrial diseases, caused by impaired OXPHOS, often present with severe neurological symptoms despite the ubiquitous expression of mitochondrial genes, such as Leigh Syndrome, optic atrophy, stroke-like episodes and peripheral neuropathy, suggesting intrinsic cell-type differences in stress responses (Gorman et al. 2016; Ng and Turnbull 2016). Understanding how mitochondrial dysfunction leads to selective neuronal loss is crucial for identifying pathways that promote neuronal resilience. Addressing this question requires direct comparative studies of how distinct neuronal subtypes respond to the same mitochondrial perturbation. While stem cell-derived neuronal models offer a powerful platform for such studies, only few have directly examined subtype-specific adaptations to mitochondrial stress under controlled conditions (Afshar-Saber et al. 2024; Lopez-Fabuel et al. 2016; Paß et al. 2021; Flønes et al. 2024).

The mitochondrial translation release factor in rescue (MTRFR) is a key quality-control factor that resolves stalled mitoribosomes translating non-stop mRNAs by catalysing peptidyl-tRNA hydrolysis, thereby releasing incomplete nascent chains (Koludarova and Battersby 2024; Ng et al. 2022). Biallelic loss-of-function mutations in *MTRFR* cause a progressive neuromuscular disorder characterised by a triad of neurological manifestations, including spasticity, peripheral neuropathy and optic atrophy, while other features such as ataxia and learning difficulties are less common (Antonicka et al. 2010; Spiegel et al. 2014). Despite its ubiquitous expression, pathology is largely confined to motor neurons, suggesting that *MTRFR* dysfunction unmasks intrinsic differences in how neuronal subtypes tolerate mitochondrial stress.

To investigate the mechanisms driving selective neuronal vulnerability, we generated a knockdown model using CRISPR interference to repress *MTRFR* expression. Isogenic iPSCs were differentiated into cortical and spinal motor neurons to examine how the same mitochondrial defect produces distinct cellular outcomes. *MTRFR* loss led to comparable reductions in mitochondrial translation and OXPHOS protein levels in both cell types but triggered divergent stress responses. Cortical neurons activated adaptive programs, such as the heat-shock response, that preserved survival and mitochondrial function, whereas motor neurons exhibited increased apoptosis and inflammatory signalling.

Pharmacological activation of the heat-shock response was sufficient to restore survival in vulnerable motor neurons, demonstrating that resilience may be programmable by mimicking intrinsic protective pathways. Together, these results establish *MTRFR* loss as a tractable model for dissecting the principles of selective neuronal vulnerability to mitochondrial dysfunction. By linking compartment-specific mitochondrial remodelling with stress adaptation, this work reframes resilience, rather than dysfunction, as the key determinant of neuronal fate under metabolic stress.

## RESULTS

### CRISPRi-mediated MTRFR knockdown disrupts mitochondrial translation in human cortical and motor neurons

To investigate how loss of *MTRFR* impacts neuronal health and mitochondrial function, we used CRISPR interference (CRISPRi) to repress *MTRFR* expression in human iPSCs (Tian et al. 2019). The catalytically inactive dCas9–KRAB repressor was targeted to the *MTRFR* transcription start site using three independent single-guide RNAs (sgRNAs), generating three knockdown lines (KD1–3) with >90% reduction in *MTRFR* transcript levels (Figure 1A). A scrambled sgRNA line and the unedited parental iPSC line were used as isogenic controls throughout all experiments.

**Figure 1.**
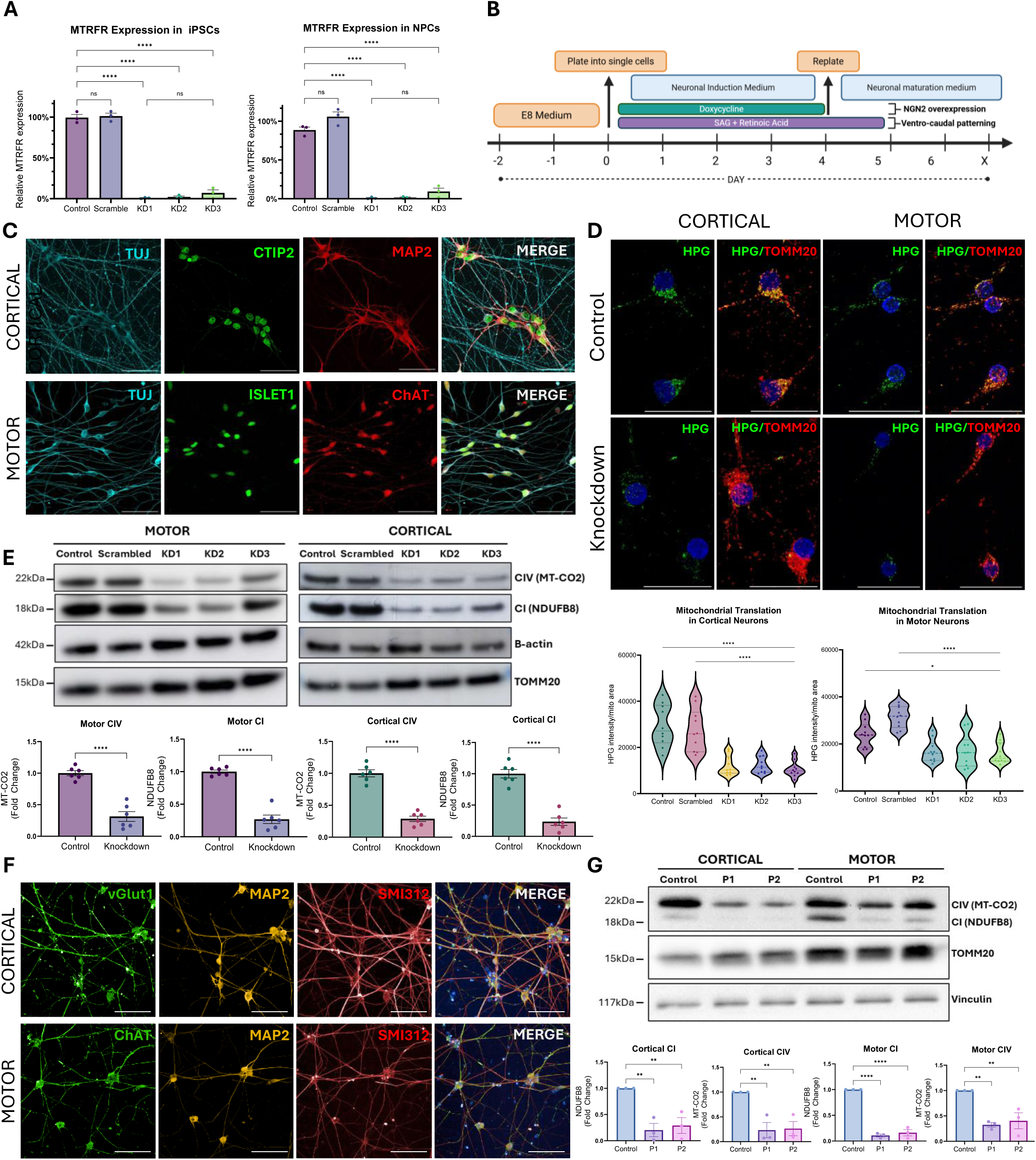
MTRFR KD impairs mitochondrial translation in cortical and motor neurons. **A** RT-qPCR analysis of *MTRFR* expression in iPSCs transduced with individual sgRNAs and in resulting NPCs, showing efficient knockdown relative to control cells. Expression was normalised to β*-actin*. Values were plotted as mean ± SEM from *n = 3* biological replicates. ****P < 0.0001, one-way ANOVA followed by Tukey’s post hoc test. **B** Schematic of the differentiation protocol used to generate cortical and motor neurons from iPSCs. Cells were replated as single cells on day 0, and neuronal induction was initiated by doxycycline-induced overexpression of *NGN2*. For motor neuron specification, SAG and retinoic acid were added for the first five days to promote ventral-caudal patterning. Cells were grown in maturation medium until day 20. **C** Immunocytochemistry for key neuronal and subtype-specific markers was used to validate the identity of differentiated neurons. General neuronal identity was confirmed by TUJ1 staining. Cortical neuron identity was supported by expression of MAP2 and CTIP2, while motor neuron differentiation was indicated by ISLET1 and ChAT expression. Scale bars, 50µm. **D** Cytosolic translation was inhibited with cycloheximide, and newly synthesised mitochondrial proteins were labelled by incorporation of the methionine analogue HPG, followed by fluorescent tagging via a click reaction with an Alexa Fluor-488 azide. Both cortical and motor neurons exhibited reduced mitochondrial translation, as indicated by decreased HPG fluorescence. Quantification of HPG signal intensity per neuron, normalised to mitochondrial area (TOMM20-positive signal), showed a significant reduction in mitochondrial translation across all three *MTRFR* KD guides in both cortical and motor neurons. Each data point in the violin plot represents an individual neuron. Quantification was performed on 10 neurons per cell line, across two independent click chemistry experiments. ****P < 0.0001, *P < 0.05, one-way ANOVA followed by Tukey’s post hoc test. **E** Western blot analysis of OXPHOS complexes revealed a significant decrease in complex I and complex IV subunits in KD cortical and motor neurons relative to controls. Band intensities were normalised to TOMM20 and quantified across *n* = 6 independent samples. Data from all KD and control lines were pooled for analysis. ****P < 0.0001, unpaired two-tailed *t*-test. **F** Immunocytochemistry of patient-derived cortical and motor neurons for key neuronal and subtype-specific markers. Matured neuronal identity was confirmed by SMI312 and MAP2. Cortical neuron identity was confirmed vGlut1 expression, while motor neuron identity was supported by positive ChAT staining. Scale bars, 50µm. **G** Western blot analysis of OXPHOS complexes revealed a significant decrease in complex I and complex IV subunits in patient-derived neurons relative to controls, paralleling the knockdown results. Band intensities were normalised to TOMM20 and quantified across *n* = 3 independent samples, *P < 0.05, unpaired two-tailed *t*-test.

Cortical neurons were derived from these iPSCs using the established NGN2 overexpression protocol (Zhang et al. 2013). To generate spinal motor neurons from the same iPSC background, we introduced ventro-caudal patterning cues for 5 days using SAG and retinoic acid during the progenitor stage as previously described (Limone et al. 2023a) (Figure 1B). Neurons were cultured for 20 days. Phase-contrast microscopy revealed the expected progression of neuronal differentiation, with iPSCs acquiring a neural progenitor-like morphology following doxycycline induction and exhibiting extensive neurite outgrowth characteristic of mature cortical and motor neurons by day 11 (Supplementary Figure S1A). Immunocytochemistry confirmed the expected neuronal identities as all lines expressed the pan-neuronal marker TUJ (βIII-tubulin). Cortical neurons were positive for the deep-layer cortical marker CTIP2 and the dendritic marker MAP2, indicating neuronal maturation. Motor neurons expressed choline acetyltransferase (ChAT) and the motor neuron transcription factor ISLET1, confirming successful differentiation (Figure 1C).

To assess whether *MTRFR* KD impairs mitochondrial translation in cortical and motor neurons we employed a click-chemistry assay (Yousefi et al. 2021). This approach provides spatial resolution of mitochondrial translation by labelling newly synthesized proteins with the methionine analogue L-Homopropargylglycine (HPG) following cytosolic translation inhibition with cycloheximide. Confocal imaging revealed a significant reduction in HPG signal intensity, normalised to mitochondrial area, in KD neurons compared to controls (Figure 1D). The reduction was consistent across all three sgRNAs and observed in both cortical and motor neurons. In untreated control cells, cytosolic HPG incorporation was readily detected and was abolished by cycloheximide treatment, validating the specificity of the assay (Supplementary Figure S1B).

To further assess the impact of *MTRFR* knockdown on mitochondrial translation, we examined the abundance of OXPHOS complex subunits. Both cortical and motor neurons exhibited a marked reduction in the mitochondrial encoded Complex IV subunits MT-CO1 (Pratt et al. 2025) and MT-CO2 (Figure 1E), a molecular hallmark of MTRFR loss of function. In parallel, the Complex I subunit NDUFB8 was also significantly decreased in both neuronal types (Figure 1E), consistent with the reduction of Complex I and IV typically observed in other more common mitochondrial translation disorders (Čunátová et al. 2024; Miller et al. 2004). In contrast, SDHB (Complex II), UQCRC2 (Complex III), and ATP5A (Complex V) levels remained unchanged when normalised to TOMM20 (Supplementary Figure S1C). Collectively, these results establish *MTRFR* knockdown neurons as a robust system to study how distinct neuronal populations adapt to mitochondrial dysfunction and to dissect mechanisms underlying selective vulnerability.

To validate these findings in a genetically relevant disease context, we examined an independent patient-derived iPSC model generated from two patients (P1 and P2) carrying biallelic pathogenic variants in *MTRFR* and a healthy control (Pyle et al. 2014). Using the same NGN2-mediated protocols applied in our CRISPRi system, patient iPSCs successfully differentiated into cortical and motor neurons expressing the expected markers, confirming that MTRFR deficiency does not impair lineage specification (Figure 1F). Notably, patient-derived cortical and motor neurons exhibited the same OXPHOS defects observed in the knockdown model, including reduced abundance of the mitochondrial-encoded Complex IV subunit MT-CO2 and decreased NDUFB8 (Figure 1G). These findings demonstrate that impaired mitochondrial translation and the resulting Complex I and IV deficiencies are shared features across both engineered and patient-derived systems, reinforcing the relevance of the CRISPRi model for mechanistic studies of MTRFR dysfunction.

### Cortical and motor neurons exhibit distinct mitochondrial responses to MTRFR-induced dysfunction

We next measured if *MTRFR* KD induces metabolic dysfunction in cortical and motor neurons. Glucose uptake was assessed by 2-deoxyglucose (2DG) incorporation using the Glucose Uptake-Glo assay and was unchanged in both cortical and motor knockdown neurons relative to controls (Figure 2A). In contrast, total intracellular ATP levels, quantified by CellTiter-Glo, were significantly reduced by 40.5% in cortical neurons and by 49.6% in motor neurons compared to controls (Figure 2B). To test whether this was accompanied by enhanced glycolytic activity, we measured lactate release using the Lactate-Glo assay. Lactate production was increased in cortical neurons by 55.7%, with a more pronounced rise observed in motor neurons (162% higher compared to control) (Figure 2C). These findings indicate a conserved energetic deficit across neuronal subtypes, characterised by a glycolytic shift coupled with a possible hypermetabolic state.

**Figure 2.**
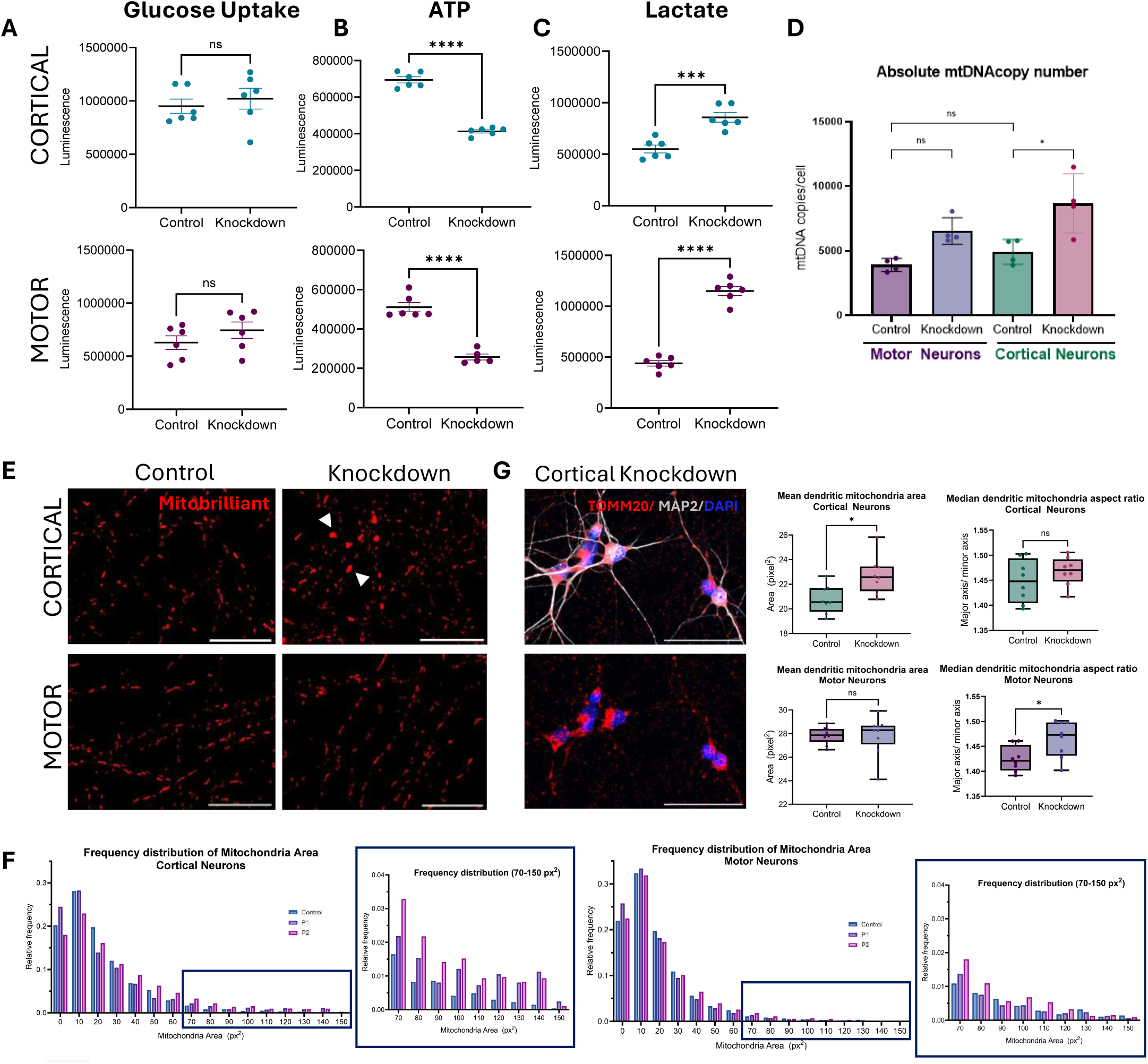
MTRFR KD induces differential mitochondrial responses in cortical and motor neurons. **A-C** Loss of MTRFR impairs mitochondrial energy production and promotes a shift toward glycolytic metabolism. **A** Glucose uptake was unchanged in both cortical and motor neurons as measured using the Glucose Uptake-Glo assay. **B** Cellular ATP levels were significantly decreased in KD neurons, as determined using the CellTiter-Glo assay. **C** ATP depletion was accompanied by a significant increase in lactate secretion, assessed using the Lactate-Glo assay. All luminescent values were normalised to cell number and plotted as mean ± SEM from *n = 6* independent samples. ****P < 0.0001, ***P < 0.001, unpaired two tailed *t*-test. **D** Digital qPCR revealed a significant increase in mtDNA copy number in cortical KD neurons, with no change observed in motor neurons. Data were plotted as mean ± SEM for n=4. **P > 0.01, unpaired two-tailed *t*-test. **E** Mitochondria were labelled with MitoBrilliant. Live imaging revealed regions of increased mitochondrial signal in cortical MTRFR KD neurons, consistent with mitochondrial enlargement. Scale bars, 25µm. **F** Enlarged mitochondria co-localised with MAP2-labelled dendrites. Quantification of dendritic mitochondria using CellProfiller showed a significant increase in average mitochondrial area in KD cortical neurons compared to controls. No significant differences were observed in motor neurons. Median aspect ratio was significantly increased in KD motor neurons. Boxplots display the mean, minimum, and maximum values with all data points shown. Eight regions of interest (ROIs) per condition were analysed, and average values per ROI were plotted. *P > 0.05, unpaired two-tailed *t*-test. **G** Enlarged mitochondria were also observed in cortical neurons differentiated from patient-derived iPSCs. Frequency distribution analysis of mitochondrial area confirmed that large mitochondria occurred at a higher rate in patient cortical neurons compared to controls, whereas no difference was apparent in motor neurons.

We next investigated whether cortical and motor neurons modulate mitochondrial DNA (mtDNA) content in response to *MTRFR* knockdown-induced mitochondrial dysfunction. Cortical neurons exhibited a significant increase in mtDNA copy number (76%) relative to controls, consistent with a compensatory response to sustain mitochondrial function. In contrast, while motor neurons showed an upward trend, the change in mtDNA content was not significant, suggesting a reduced capacity to compensate for the mitochondrial translation defect (Figure 2D).

To determine whether these molecular alterations were accompanied by structural remodelling, we assessed mitochondrial morphology in cortical and motor neurons using MitoBrilliant staining. Cortical KD neurons exhibited regions containing enlarged mitochondria within neuronal projections that were not present in motor KD neurons (Figure 2E). A similar increase in enlarged mitochondria was observed in patient-derived cortical neurons, and frequency distribution analysis confirmed that large mitochondria occurred at a higher rate in patient cells compared to controls, whereas no difference was apparent in motor neurons (Figure 2F, Supplementary Figure S2B). To determine where these enlarged mitochondria localise within neurons, we performed immunofluorescence analysis, which showed that they were predominantly present within MAP2-positive dendritic regions. Notably, approximately 5% of dendritic mitochondria were enlarged, with an area exceeding 3.5µm², resulting in a significant increase in mean dendritic mitochondrial area (Figure 2G). In contrast, mitochondrial area in motor neurons remained unchanged, although aspect ratio appeared significantly increased in motor KD neurons, suggesting more subtle alterations in mitochondrial shape.

Together, these findings reveal that while cortical and motor neurons share comparable metabolic defects, their mitochondrial responses diverge, with motor neurons displaying a reduced adaptive capacity to MTRFR-induced dysfunction.

### Enlarged mitochondria in cortical neurons act as local metabolic hubs

To examine the structure of the enlarged mitochondria seen only in MTRFR KD cortical neurons we performed transmission electron microscopy (TEM). TEM images confirmed that swollen mitochondria are present in cortical neurites either as single mitochondrion or as small clusters in discrete regions. These structures contained electron-dense foci, likely corresponding to nucleoid accumulation, while maintaining intact cristae (Figure 3A). Consistent with the TEM observations, these enlarged mitochondria were strongly positive for TFAM, confirming the presence of nucleoid-rich domains (Figure 3B, Supplementary figure S2A). In addition, DNA staining further verified that these mitochondria contained densely packed mitochondrial DNA (Figure 3C).

**Figure 3.**
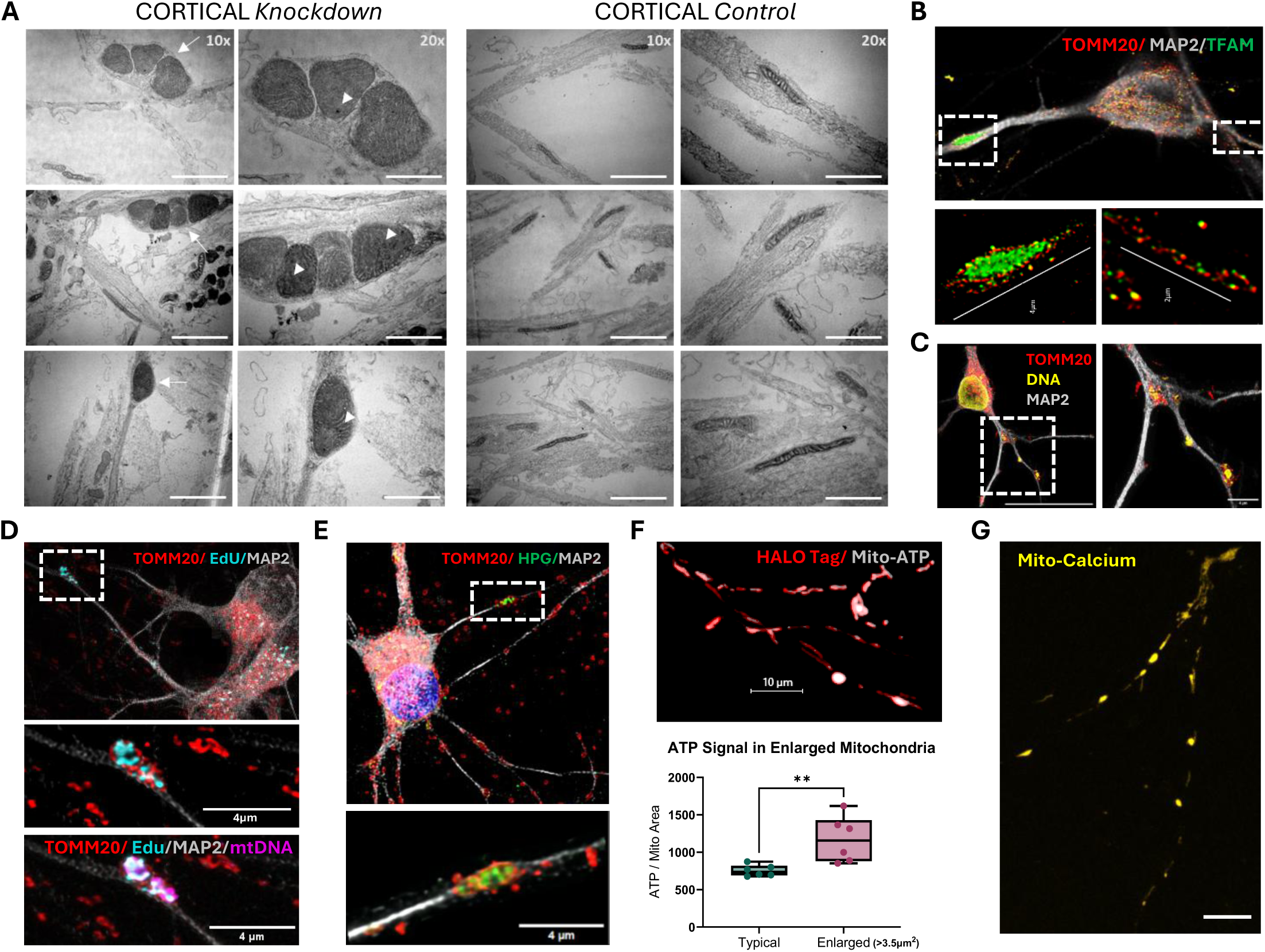
Enlarged mitochondria in cortical neurons exhibit localised mtDNA Replication and active metabolism. **A** Transmission electron microscopy (TEM) of cortical neurons revealed that enlarged mitochondria can present either isolated single organelles or as multiple clustered mitochondria within dendrites, often distorting the local dendritic architecture (arrow). Electron-dense regions suggestive of nucleoid accumulation were also observed (arrowhead). In contrast, mitochondria in control neurons displayed typical tubular morphology. Scale bars, 2µm (10x magnification) and 1µm (20x magnification). **B** STED microscopy of TFAM, TOMM20 and MAP2, showed single enlarged mitochondria in dendrites of cortical neurons filled with TFAM signal, suggesting nucleoid accumulation. Scale bars 4 µm, 2µm. **C** Co-labelling of MtDNA, TOMM20 and MAP2 in knockdown cortical neurons confirmed the presence of nucleoid clusters within enlarged dendritic mitochondria. Scale bars, 25µm (overview) and 4µm (inset). **D** EdU click-chemistry replication assay showed active mtDNA replication within enlarged mitochondria, co-localizing with clustered nucleoids. Scale bars 4 µm. **E** HPG click-chemistry translation assay showed ongoing mtDNA translation in enlarged mitochondria, despite an overall reduction in global mitochondrial translation levels. Scale bars, 4 µm. **F** A genetically encoded mitochondrial ATP sensor indicated increased ATP signal in enlarged mitochondria. Quantification of ATP integrated density normalised to mitochondria area showed significantly higher ATP levels in enlarged mitochondria compared to typical mitochondria within the same cell. Mitochondria were classified as enlarged if area >3.5µm². ATP values were normalised per mitochondrion, then averaged per cell for n= 6 cells. Boxplots display the mean, minimum, and maximum values with all data points shown. Scale bars, 10µm and 25µm. **G** Mitochondrial calcium imaging using a genetically encoded sensor shows bright calcium signal in enlarged mitochondria, indicating calcium accumulation. Scale bar, 10µm.

We next examined whether these nucleoid-rich enlarged mitochondria were functionally active. The enlarged mitochondria coincided with sites of active mtDNA replication, evidenced by increased EdU incorporation (Figure 3D). The click-chemistry translation assay also revealed locally elevated HPG signal within these structures, despite the global reduction in mitochondrial translation (Figure 3E). Similarly, live-cell imaging using a mitochondrially targeted ATP sensor (Marvin et al., 2024) showed higher ATP signal within enlarged mitochondria compared to adjacent regions (Figure 3F), indicating local metabolic activity. Increased mitochondrial matrix calcium, consistent with enhanced TCA flux and ATP synthesis (Glancy et al. 2013; Phillips et al. 2012), further supported that these large mitochondria are functional and active (Figure 3G).

Because enlarged mitochondria can potentially interfere with organelle trafficking, we investigated whether mitochondrial motility was affected. Kymograph-based analysis revealed no significant difference in mitochondrial movement between MTRFR knockdown and control cortical neurons (Supplementary Figure S2C), indicating that these structures did not lead to trafficking defects in the short term.

Together, these findings indicate that enlarged dendritic mitochondria seen only in cortical neurons act as localised hubs that sustain focal bioenergetic activity despite global mitochondrial dysfunction and represents a potential compensatory mechanism, supporting survival of cortical, but not motor neurons.

### Cortical resilience to MTRFR loss is associated with mitochondrial plasticity and adaptive stress responses

To identify molecular pathways underlying differential neuronal resilience, we performed bulk RNA-seq on cortical and motor neurons after *MTRFR* knockdown. Principal component analysis revealed clear clustering by cell type (Supplementary Figure S3A). Differential expression analysis between control and KD neurons ran individually for each subtype identified 1,558 downregulated and 436 upregulated genes in cortical neurons, and 160 downregulated and 373 upregulated genes in motor neurons (Figure 4A). Moreover, gene set enrichment analysis (GSEA) revealed distinct adaptive programs across subtypes.

**Figure 4.**
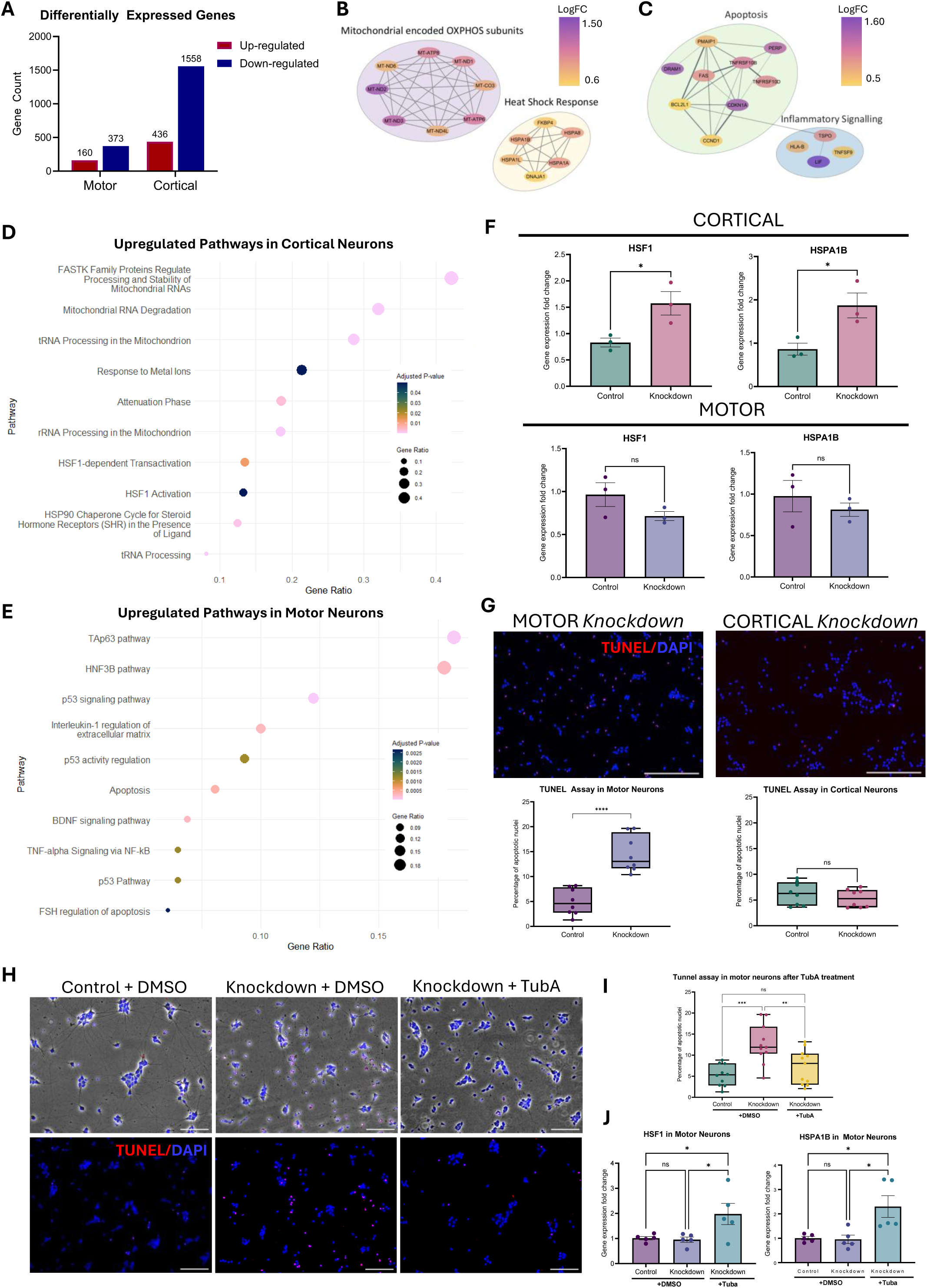
Differential stress-adaptive programs underlie cortical vs motor neuron vulnerability to MTRFR loss and predict pharmacological rescue. A-G Cortical and motor neurons activate distinct transcriptomic programs and exhibit differential stress-response capacities. **A** Bar plot of differentially expressed genes for both cortical and motor neurons. A gene was considered differentially expressed at FDR < 0.05 and |logFC| > 0.5. **B-C** Protein-protein interaction network showing key genes significantly upregulated after MTRFR KD, nodes represent significantly upregulated genes (FDR < 0.05), edges represent known or predicted functional associations, and node colour corresponds to log_2_ fold-change. **B** Cortical neurons exhibit induction of mitochondrial gene expression and HSP70-mediated stress-adaptive pathway. **C** Motor neurons show upregulation of pro-apoptotic and inflammation-related transcripts. **D-E** Gene set enrichment analysis (GSEA) of differentially expressed genes in KD cortical and motor neurons. The top 10 enriched pathways, ranked by adjusted p-value, were plotted. Dot colour represents the adjusted p-value, and dot size corresponds to the gene ratio for each pathway. **D** GSEA shows upregulation of adaptive stress response pathways in cortical neurons, including the heat shock response and ATP synthesis-related programs. **E** In motor neurons, apoptotic and inflammatory pathways were significantly upregulated in response to *MTRFR* KD. **F** RT-qPCR analysis showed that *MTRFR* KD induced expression of the heat shock response (HSR) transcription factor HSF1 and HSP70 (HSPA1B) in cortical but not motor neurons. Expression was normalised to β*-actin*. Data were plotted as mean ± SEM for n=3. *P < 0.05, unpaired two tailed *t*-test. **G** TUNEL assay revealed a significantly higher percentage of apoptotic nuclei in MTRFR KD motor neurons compared to controls, while no change was observed in cortical neurons. Boxplots display the mean, minimum, and maximum values for n=8 regions of interest (ROI), with each data point representing the average per ROI. ****P < 0.0001, unpaired two tailed *t*-test. Scale bars, 200 µm. **H-J Pharmacological activation of the heat-shock response rescues motor neuron survival. H-I** Treatment with the HDAC6 inhibitor Tubastatin A (TubA) for 24 hours significantly reduced the percentage of TUNEL-positive nuclei in motor neurons, restoring apoptosis levels to control. Scale bars, 100 µm. Boxplots show the mean, minimum, and maximum values for n=10 regions of interest (ROIs), with each data point representing the average per ROI. ***P < 0.001, **P < 0.01, one-way ANOVA followed by Tukey’s post hoc test. **J** TubA treatment significantly increased expression of HSR-related genes in motor neurons, suggesting the observed rescue is at least partially mediated by HSR activation. Expression was normalised to β*-actin*. *P < 0.05, one-way ANOVA with Tukey’s post hoc test.

Cortical neurons showed strong induction of stress-adaptive pathways, including mitochondrial gene expression and protein homeostasis (Figure 4B, D). Over half of the mitochondrially encoded transcripts, including *MT-ND6*, *MT-ATP6*, *MT-CO3*, *MT-ND2*, *MT-ND3*, *MT-ATP8*, and *MT-ND1*, were selectively increased, consistent with GSEA enrichment of mitochondrial translation and OXPHOS-related terms. Additionally, members of the HSP70 family (*HSPA8*, *HSPA1L*, *HSPA1B*, *HSPA1A*) and the heat-shock response (HSR) regulator *HSF1* were also elevated, suggesting HSR activation. Of note, HSP70 induction is known to exert neuroprotective effects across models of neurodegeneration (Turturici et al. 2011; Paul and Mahanta 2014). Together, these results suggest that cortical neurons engage a protective transcriptional response to MTRFR loss, potentially contributing to their relative resilience.

In contrast, knockdown motor neurons displayed a transcriptional signature dominated by apoptosis and inflammation related pathways (Figure 4C, E). Upregulated apoptosis-related genes included *FAS*, *TNFRSF10B*, *PMAIP1*, *BCL2L11* and *DRAM1*, consistent with activation of both death receptor and mitochondrial apoptotic pathways. In parallel, increased expression of inflammatory and immune-response genes such as *LIF*, *TNFSF9*, *HLA-B* and *TSPO* suggested activation of innate immune and interferon-related signalling. Nevertheless, phosphorylated STING and cleaved caspase-1 were not increased in these cells (Supplementary Figure S3C), suggesting inflammation is secondary to apoptosis rather than its cause. This pattern is consistent with many models of ALS in which motor neuron degeneration is thought to be initiated by intrinsic defects, with glial activation arising later and subsequently contributing to disease progression (Boillée et al. 2006; Brites and Vaz 2014).

### Pharmacological enhancement of the heat-shock response rescues motor neuron survival

To validate the transcriptional evidence for selective activation of the HSR, we examined expression of key HSR components in MTRFR KD cortical and motor neurons. RT-qPCR confirmed increased expression of *HSF1*, the master regulator of the HSR, and its downstream target *HSPA1B* in *MTRFR* knockdown cortical neurons, whereas no change was detected in motor neurons (Figure 4F). These results demonstrate that the HSR is selectively induced in cortical neurons, consistent with their transcriptional profile. The combined induction of mitochondrial and cytoprotective programs suggests that cortical neurons mount a coordinated adaptive response that integrates mitochondrial remodelling with HSP70 induction to maintain cellular energy balance and enhance their resilience.

To assess whether these molecular differences correlate with cell vulnerability, apoptosis in cortical and motor neurons was assessed using the terminal deoxynucleotidyl transferase dUTP nick end labelling (TUNEL) assay. At day 15, the proportion of TUNEL-positive nuclei were significantly increased in *MTRFR* knockdown motor neurons compared to controls, while cortical neurons remained unaffected (Figure 4G).

## DISCUSSION

Selective neuronal vulnerability is a hallmark of many mitochondrial and neurodegenerative disorders, yet the mechanisms that determine why certain neuronal populations are more susceptible while others remain resilient are poorly understood. Here, using the loss of *MTRFR* as a model of mitochondrial dysfunction, we show that neuronal resilience depends not only on the degree of metabolic impairment but on intrinsic differences in mitochondrial plasticity and stress-adaptive capacity between neuronal cell types. Despite comparable reductions in mitochondrial translation and OXPHOS complexes, as well as similar metabolic rewiring, motor neurons were more susceptible to *MTRFR* loss, exhibiting increased apoptosis, whereas cortical neurons survived by engaging adaptive programs. These findings indicate that selective vulnerability arises from cell-type-specific thresholds for adaptation rather than conserved mitochondrial failure. Replication of the findings in two independent in vitro models, including patient-derived iPSCs differentiated into cortical and motor neurons, confirms the robustness of these cell-type-specific responses.

Cortical neurons compensated for impaired mitochondrial translation by upregulating mitochondrially encoded transcripts and increasing mtDNA copy number, consistent with enhanced mitochondrial gene expression. Morphological and ultrastructural analyses revealed enlarged, metabolically active mitochondria within dendrites, characterised by clustered nucleoids, increased mtDNA replication and translation, and elevated local ATP production and calcium buffering, features indicative of localised mitochondrial plasticity. These metabolically enriched regions likely act as compartmentalised energy reservoirs that allow cortical neurons to buffer mitochondrial dysfunction. In contrast, motor neurons failed to exhibit such structural or metabolic remodelling, highlighting fundamental differences in how neuronal subtypes maintain mitochondrial homeostasis under stress.

The dendritic localisation of these adaptive mitochondria likely reflects cell-type-specific differences in subcellular organisation. In cortical neurons, mitochondria in dendrites are spatially anchored through the cytoskeleton, forming defined compartments that support local energy demands and synaptic activity (Virga et al. 2024a). In response to mitochondrial stress, these dendritic mitochondria may increase their metabolic output, leading to mtDNA clustering and mitochondrial swelling. By contrast, motor neurons rely on simpler dendritic architecture and long-range mitochondrial transport, with synaptic inputs located more proximally (Zaninello and Bean 2023; Virga et al. 2024b; Rangaraju et al. 2014). This structural organisation may restrict their capacity for local metabolic compensation, contributing to their selective vulnerability. Thus, architectural asymmetries likely define the limits of mitochondrial adaptation across neuronal types.

Pharmacological induction of the HSR significantly improved survival of *MTRFR*-deficient motor neurons, confirming that engaging this pathway is sufficient to mitigate apoptosis even without directly targeting the underlying translation defect. Our results support a model in which selective neuronal vulnerability arises from differences in mitochondrial plasticity and stress-adaptive capacity rather than the severity of the primary metabolic impairment. Cortical neurons integrate these mechanisms to maintain function and energy balance, whereas motor neurons, constrained by architecture and limited stress responsiveness, are less able to compensate and therefore degenerate more readily. Enhancing adaptive mechanisms, such as through pharmacological HSR activation, represents a promising strategy to promote neuronal survival in mitochondrial and potentially other neurodegenerative diseases.

The observation that stress-pathway activation can rescue motor neuron survival highlights a therapeutic avenue complementary to gene replacement approaches. While AAV-mediated expression of wild-type *MTRFR* successfully restored OXPHOS protein levels in knockdown neurons (Pratt et al. 2025), boosting endogenous adaptive responses provides an additional route to protection that may extend to a wider range of mitochondrial disorders. Future work should examine the long-term impact of HDAC6 inhibition on mitochondrial function and neuronal integrity and test whether similar stress adaptive mechanisms operate in other vulnerable populations, such as retinal ganglion cells, as optic neuropathy is also a key feature of *MTRFR*-related disease.

Together, these findings establish *MTRFR* loss as a powerful model for dissecting the mechanisms driving selective neuronal vulnerability in mitochondrial dysfunction. They demonstrate that resilience and vulnerability are governed not only by metabolic competence but by the ability to coordinate compartment-specific mitochondrial remodelling with stress signalling. By identifying dendritic mitochondrial plasticity and HSP70 induction as hallmarks of neuronal adaptation, our work highlights therapeutic opportunities that move beyond correcting bioenergetic defects toward strategies that focus on enhancing intrinsic cellular resilience.

## ACKNOWLEDGMENTS

We would like to thank Huw Naylor, Dr. Andreas Bruckbauer, Dr. Fadwa Joud, Heather Zecchini at the Light Microscopy Core Facility from the Cancer Research UK Cambridge Institute for their assistance with microscopy and data analysis. We would also like to thank the Genomics facility from the Cambridge Stem Cell Institute for the RNA sequencing library preparation and Louis Elfari at the Electron Microscopy Facility from the Cambridge Stem Cell Institute for the training and guidance. We thank Michael Ward for the gift of the CRISPRi-NGN2-iPSC line. We would like to thank the Human Pluripotent Stem Cell Facility (HPSCF) (RRID:SCR_027437) at the Ottawa Hospital Research Institute for the reprogramming of the patient iPSCs. Dr. James Ellis from Sick Kids generously donated the PGPC-3 control line to the HPSCF for use in this project. Additionally, we would like to thank Dr. Stephen Baird at CHEO Research Institute for image acquisition using the OPERA Phenix system.

## Author Contributions

Conceptualisation: MZM, DH, RH; Methodology: MZM, JK, ER; Investigation: MZM, KO, NM; Formal Analysis: MZM, KO, OP; Visualisation: MZM; Manuscript Writing: MZM; Manuscript Reviewing: DH, RH; Supervision: DH, RH, SS, HL; Funding Acquisition: RH, HL

## SUPPLEMENTARY MATERIAL

**Supplementary Figure 1. Differentiation of neurons and OXPHOS. A** Representative bright-field images show key stages of the differentiation process. Three days after initial *NGN2* induction, neurite outgrowth was visible. Cells were replated at day 4, and by day 11 cells exhibited mature neuronal morphology. Scale bars, 250µm. **B** In the absence of cycloheximide treatment, HPG signal revealed prominent regions of cytosolic translation within the nucleus, distinct from mitochondrial labelling (arrows). **C** Western blot analysis of OXPHOS complexes show no significant difference in complex II, complex III, and complex IV subunits in KD cortical and motor neurons relative to controls. Band intensities were normalised to TOMM20 and quantified across *n* = 6 independent samples. Data from all KD and control lines were pooled for analysis. ****P < 0.0001, unpaired two-tailed *t*-test.

**Supplementary Figure 2. Nucleoid clustering and axonal transport of mitochondria in MTRFR knock down and patient-derived MTRFR mutant neurons. A** Immunocytochemistry of cortical and motor MTRFR knock down neurons revealed increased TFAM signal selectively in cortical KD neurons, consistent with nucleoid clustering. Scale bars, 5µm. **B** Mitochondria were labelled with Mitobrilliant in patient-derived cortical and motor neurons. Scale bars, 50µm. **C** Kymograph analysis from live imaging of axonal transport in cortical and motor neurons revealed no significant differences in the percentage of motile mitochondria, average velocity, or the number of mitochondria per 50µm axon segment between control and KD conditions. Mitochondria were classified as motile if their average frame-to-frame velocity exceeded 0.2µm/sec. Boxplots display the mean, minimum, and maximum values with all data points shown. n = 25 axons for cortical neurons and n = 15 axons for motor neurons. Statistical comparisons were performed using unpaired two-tailed t-tests.

**Supplementary Figure 3. Gene expression and immunoblotting for cellular responses in MTRFR knock down neurons. A** PCA of VST-transformed gene expression data showed clear separation of samples by cell type, with control and KD samples showing distinct clustering within each neuronal subtype. **B** Boxplots of VST-transformed counts for each sample displayed uniform distribution after normalisation. **C** Western blot analysis of phosphorylated STING/STING and cleaved Caspase-1/total Caspase-1 in motor neurons showed no significant differences between KD and control, suggesting that increased inflammation is a consequence rather than a driver of apoptosis. Quantification of band intensity ratios plotted as mean ± SEM for n = 4. **D** Motor neurons were treated with the proteasome inhibitor Mg132, and protein lysates were analysed by Western blot using an anti-ubiquitin antibody. MG132 treatment led to the expected high molecular weight smear, reflecting accumulation of polyubiquitinated proteins. However, the extent of ubiquitin accumulation was comparable between control and knockdown cells, suggesting there is no increased proteotoxic stress in knockdown neurons. The integrated density of the smear above 70kDa was measured and normalised to β-actin band intensity. Data were plotted as mean ± SEM for n = 3, *P < 0.05, one-way ANOVA followed by Tukey’s post hoc test.

**Supplementary Table S1.** Single guide RNAs targeting transcription start site of MTRFR

**Supplementary Table S2.** *List of primers used*

## METHODS

### RESOURCE AVAILABILITY

#### Lead contact

Further information and requests for resources and reagents should be directed to and will be fulfilled by the lead contact, Rita Horvath.

#### Materials availability

This study did not generate new unique reagents. Plasmids used in this study are available from Addgene as indicated. Lentiviral sgRNA constructs are available upon request to the lead contact, subject to institutional MTAs.

#### Data and code availability

- **Data:** Bulk RNA-seq data have been deposited at the Gene Expression Omnibus (GEO), which is hosted by the National Center for Biotechnology Information (NCBI) (in progress).
- **Code:** This paper does not report original code. Analysis scripts used for RNA-seq and image processing are available from the lead contact upon request.
- **Additional information:** Any additional information required to reanalyse the data reported in this paper is available from the lead contact.

### EXPERIMENTAL MODEL AND SUBJECT DETAILS

#### Human iPSC culture and maintenance

Patient fibroblasts were obtained from a sibling pair, one female (Patient 1) and one male (Patient 2) both presenting with homozygous *MTRFR* mutations (c.96_99dupATCC; p.Pro34Ilefs*25). Patient iPSC lines were generated by the Human Pluripotent Stem Cell Facility at the Ottawa Hospital Research Institute from patient fibroblasts using episomal reprogramming as described below.

Human iPSC lines were maintained under feeder-free conditions on Geltrex- or Matrigel-coated plates (Thermo Fisher Scientific, Corning) and cultured in either mTeSR Plus (STEMCELL Technologies) or Essential 8 (Thermo Fisher Scientific) with daily medium changes. Colonies were passaged every 6–7 days using ReLeSR (STEMCELL Technologies) or EDTA. For cryopreservation, cells were frozen in 90% mTeSR Plus/10% DMSO. Lines stably expressing dCas9-KRAB and with doxycycline-inducible NGN2 (a gift from Dr. Evan Reid) were used as described below. The PGPC-3 control line was a gift Dr. James Ellis at Sick Kids (Hildebrandt et al).

### METHOD DETAILS

#### CRISPRi-mediated knockdown of *MTRFR*

iPSCs stably expressing dCas9-KRAB were transduced with lentiviral sgRNAs targeting the *MTRFR* transcription start site (8 independent guides) or a scrambled control (CRISPick; sequences in Table SX). Oligos were phosphorylated/annealed (T4 ligase buffer, T4 PNK) and ligated into a BsmBI-digested lentiGuide-puro backbone (pKLV-U6gRNA(BbsI)-PGKpuro2ABFP; Addgene #50946). Vector-only ligation served as negative control.

Ligations were transformed into Stbl3 competent cells and 2–3 colonies per sgRNA were mini-prepped (Qiagen) and Sanger-sequenced. Lentivirus was produced in HEK293T cells using a second-generation system (p8.91 and pMD.G; Addgene #187440 and #187441) with TransIT-293 in Opti-MEM. Medium was replaced with Essential 8 after 24 h. At 48h post-transfection, the viral supernatant was collected, filtered (0.45 µm), and snap-frozen.

For transduction, iPSCs were plated as single-cells and incubated with viral supernatant (1:1 in E8) for 24 h, recovered overnight in fresh medium, and selected with puromycin starting 48 h post-transduction. Surviving pools were expanded and knockdown efficiency was assessed by RT-qPCR for *MTRFR*. pKLV-U6gRNA(BbsI)-PGKpuro2ABFP was a gift from Kosuke Yusa (Addgene plasmid # 50946; http://n2t.net/addgene:50946; RRID:Addgene_50946). pMDG and p8.91 were a gift from Simon Davis (Addgene plasmids # 187440, # 187441; http://n2t.net/addgene:187440, http://n2t.net/addgene:187441; RRID:Addgene_187440, RRID:Addgene_187441).

#### Reprogramming of patient fibroblasts into iPSCs

Patient-derived iPSCs were generated through episomal reprogramming of fibroblasts following the Okita et al. (2011) protocol. In brief, transcription factors Oct3/4 (pCXLE-hOct3/4 shp53-F), SOX2 (pCXLE-hSK), KLF4 (pCXLE-hSK), L-MYC (pCXLE-hUL)and LIN28 (pCXLE-hUL) as well as a shRNA against p53 (pCXLE-hOct3/4 shp53-F) were electroporated into human fibroblasts using the P2 Primary Cell 4D-Nucleofector X Kit L (Lonza, V4XP-2024) and shock code EN-150. 72 h after electroporation media was changed to E7 with 0.25mM sodium butyrate, media was changed daily for 7 days. iPSC-like colonies were reverse picked as they appeared, and media was switched to mTeSR Plus. Pluripotency was confirmed via immunofluorescent staining for Nanog, Oct3/4 and SOX2, as well as trilineage differentiation (RRID:SCR_027437). Clones have a normal karyotype (WiCell Laboratory) and tested negative for mycoplasma (RRID:SCR_027437). Reprogramming vectors pCXLE-hOct3/4 shp53-F (Addgene #27077), pCXLE-hSK (Addgene #27078), pCXLE-hUL (Addgene #27080) and pCXLE-eGPP (Addgene #27082) were gifts from Shinya Yamanaka.

#### Generation of patient Ngn2 iPSC lines

A lentivirus expressing Ngn2 (pTet-O-Ngn2-puro; Addgene: #52047) and a doxycycline-inducible promoter (FUdeltaGW-rtTA; Addgene: #19780) was generated using the pPACKH1 HIV Lentivector Packaging Kit (System Bio) following manufacturer’s instructions using HEK293 cells. The virus was precipitated with the PEG-it Virus Precipitation Solution (System Biosciences). For transduction, iPSCs were plated in small colonies into 12-well plates on Matrigel. 24h after reseeding, cells were incubated with 8μg/mL of polybrene, 10μM Y-27632 (ROCK inhibitor) and 20μL of the viral preparation for 16h. Cells were expanded and cryopreserved as described above. Efficiency was tested by treating cells with 4μg/mL doxycycline for 24h, before selecting the cells with 2μg/mL of puromycin. FUdeltaGW-rtTA was a gift from Konrad Hochedlinger (Addgene plasmid # 19780; http://n2t.net/addgene:19780; RRID:Addgene_19780). pTet-O-Ngn2-puro was a gift from Marius Wernig (Addgene plasmid # 52047; http://n2t.net/addgene:52047; RRID:Addgene_52047)

#### Transcription factor-mediated neuronal differentiation

iPSCs stably expressing a doxycycline-inducible NGN2 transgene were used for direct neuronal differentiation of genetically modified cell lines as follows:

**Cortical neurons:** iPSCs were dissociated with Accutase (6 min, 37°C) and plated on Geltrex-coated plates in neuronal induction (NI) medium (DMEM/F12, N2, MEM-NEAA) with 10 µM Y-27632 and 2 ng/mL doxycycline. NI + doxycycline was refreshed daily for 3 days. On day 4, progenitors were dissociated with Accutase and replated on PEI-treated, Geltrex-coated plates in BrainPhys medium supplemented with SM1 (with vitamin A), BDNF (10 ng/mL) and NT3 (10 ng/mL). Media changes were performed every 3-4 days until collection.

**Lower motor neurons:** During neuronal induction, progenitors were patterned with retinoic acid (RA, 1 µM), SAG (1 µM), SB431542 (10μM), and LDN193189 (200nM) with media changes performed daily. On day 4, cells were replated in BrainPhys with SM1 (with vitaminA) plus BDNF (10 ng/mL), GDNF (10 ng/mL) and CNTF (10 ng/mL). Media changes were performed every 3-4 days until collection.

Patient-derived iPSC neuronal differentiation was carried out as follows:

**Cortical neurons:** Cortical neurons were generated as described in Wang et al. (2023). iPSCs were dissociated with EDTA (3min, room temperature) and plated on Matrigel-coated plates in small colonies. When plates reached 80% confluency the media was replaced with mTeSR Plus with 4μg/mL doxycycline on Day 0. On Days 1 & 2 media was refreshed with neuronal induction media (NI) (N2, GlutaMAX and MEM-NEAA in DMEM-F12) supplemented with NT3 (10ng/mL), BDNF (10ng/mL), doxycycline (4μg/mL) and puromycin (2μg/mL). From days 3-5 the media was refreshed every other day with cortical maturation media (CMM) (B27 and GlutaMAX in Neurobasal) supplemented with NT3 (10ng/mL), BDNF (10ng/mL), Ara-C (2μM), doxycycline (4μg/mL) and puromycin (2μg/mL). On Day 6, the neural cells were dissociated with Accutase and replated on poly-D-lysine (PDL) treated and laminin-coated (15μg/mL) Ibidi 96-well plates. A half media change was performed with CMM supplemented with NT3 (10ng/mL) and BDNF (10ng/mL) every 2-3 days until endpoint. Cells collected for protein, RNA and DNA were not replated on Day 6.

**Lower motor neurons:** Lowe motor neurons were generated as described in Limone et al. (2023b). iPSCs were dissociated with EDTA (3min, room temperature) and plated on Matrigel-coated plates in small colonies. When plates reached 80% confluency the media was replaced with NI supplemented with SB431542 (10μM), SAG (2μM), LDN193189 (200nM), retinoic acid (2μM) and doxycycline (4μg/mL) on Day 1. On Day 2 & 3 puromycin selection was performed (2μg/mL). On days 4-6, the media was changed daily with Motor Maturation Media (MMM) (MEM-NEAA, GlutaMAX, N2 and B27 in Neurobasal) supplemented with BDNF (10ng/mL), GDNF (10ng/mL), CNTF (10ng/mL), SAG (1μM), retinoic acid (1μM), doxycycline (4μg/mL) and puromycin (2μg/mL). On Day 7, cells were dissociated with Accutase and replated on poly-D-lysine (PDL) treated and laminin-coated (15μg/mL) Ibidi 96-well plates. Half media changes were performed every 2-3 days with MMM supplemented with BDNF (10ng/mL), GDNF (10ng/mL) and CNTF (10ng/mL) until endpoint. Cells collected for protein, RNA and DNA were not replated on Day 6.

#### Protein extraction and immunoblotting

Neurons were washed in PBS and lysed in SDS lysis buffer or RIPA buffer with protease/phosphatase inhibitors and lysates were sonicated. Ten to twenty micrograms of protein were resolved by SDS-PAGE and transferred to a PVDF membrane (dry transfer or semi-dry transfer). Membranes were blocked (5% milk/TBST or Intercept Blocking Buffer), incubated with primary antibodies overnight (4°C), then with HRP-conjugated secondary antibodies (1 h, RT). Membranes were imaged by chemiluminescence and band intensities quantified in ImageJ/Fiji. Signals were normalized to TOMM20 (mitochondrial proteins) or β-actin (nuclear-encoded proteins).

#### mtDNA copy number quantification

Genomic DNA was extracted with a DNeasy Blood & Tissue kit (Qiagen).

**Relative copy number (qPCR):** Quantified as previously described by Rooney et al. (2015) using a multiplex TaqMan assay (Bio-Rad CFX96) targeting MT-ND1 (mtDNA) and B2M (nuclear). Cycling: 95°C 3 min, 40 × (95°C 10 s; 62.5°C 1 min), hold at 4°C. Both targets were quantified on the same plate and samples were run in triplicates. Relative mtDNA = ΔCt(nuclear – mitochondrial).

**Absolute copies per cell (ddPCR):** Quantified following the protocol published by O’Hara et al. (2019) on a Bio-Rad QX200 system, targeting MT-ND4 (mtDNA) and RNaseP (nuclear). DNA input: 0.2 ng (mtDNA) and 8 ng (nDNA). Cycling: 95°C 10 s, 40×(94°C 30 s; 60°C 60 s), hold at 4°C. Droplets were read in QuantaSoft and absolute copies per cell were calculated from positive droplet counts for MT-ND4 and RNaseP (nuclear genome copies ×2). Primers/probes are listed in Table S2.

#### Reverse-transcription quantitative PCR (RT-qPCR)

RNA (RNeasy, Qiagen) was reverse transcribed with iScript gDNA Clear cDNA kit (Bio-Rad). Quantitative PCR was performed using the SYBR Green Supermix from Bio-rad on a CFX96 system: 95°C 3 min, 40 × (95°C 10 s, 60°C 30 s), melt curve 65–95°C (0.5°C/5 s). Expression was calculated by 2^−ΔΔCt relative to β-actin. Primer sequences are provided in Table S3.

#### Immunocytochemistry and TUNEL

Cells were fixed in 4% PFA (12 min, RT), permeabilized with 0.3% Triton X-100 (15 min, RT), and blocked in 1% BSA/PBS (1 h, RT). Primary antibodies (Table S4) were applied overnight (4°C) and Alexa-conjugated secondaries were applied for 1 h (RT). Nuclei were counterstained with DAPI (0.2 µg/mL) and samples mounted in ProLong Gold or ProLong Diamond.

Apoptosis was assessed by TUNEL per manufacturer’s instructions. Following fixation/permeabilization, samples were incubated in TUNEL equilibration buffer, then TUNEL reaction buffer mixed with TdT Enzyme (1 h, 37°C). Samples were subsequently co-stained with primary and secondary antibodies as described above.

#### Click-chemistry assays for mitochondrial translation and mtDNA replication

**Mitochondrial translation:** live neurons were incubated in methionine-free medium (30 min), treated with cycloheximide (20 min) to inhibit cytosolic translation, then labelled with the methionine analogue HPG (1 h, 37°C).

**mtDNA replication:** neurons were incubated with EdU (10 µM) overnight. Ethidium bromide co-incubation served as a negative control.

For both assays, cells were treated with 0.015% digitonin in HEPES buffer (2 min, on ice), then fixed, permeabilized, and blocked (as above). Click reactions were performed using the Click-iT Cell Reaction Buffer Kit (Thermo Fisher) with Alexa Fluor 488-azide (3 µM, 30 min, RT), followed by immunostaining and mounting.

#### Live-cell labelling and lipofection

For mitochondrial Ca²⁺ imaging, cells were transfected at low efficiency with mito-R-GECO1 (Addgene #46021) using Stem Lipofectamine (40 min, 37°C). Imaging was carried out ≥48 h post-transfection. For mitochondrial ATP, cells were transfected with mito-iATPSnFR2-HaloTag (Addgene #209722) and labelled with a HaloTag ligand prior to imaging.

For mitochondrial transport, neurons were either stained with MitoBrilliant (75 nM, 40 min) or transfected with PGK-4xMito-mEmerald (Addgene #200430) using Stem Lipofectamine for sparse labelling.

#### Image acquisition and processing

Labelled cells were imaged using 20x, 40x, 60x, and 100x objectives. Fixed samples were imaged on a Leica Stellaris 8 confocal microscope, live imaging was done using an Andor Dragonfly 200 spinning-disk system, and super-resolution imaging was done on a Leica SP8-STED. Fixed-cell images were acquired at 1024×1024 with Z-steps matched to sample thickness. STED depletion laser was set to 85%. For click-chemistry assays, LIGHTNING (auto-deconvolution) at 1.28× zoom was used. Laser power and detector gain were held constant within experiments.

Live imaging was performed in a CO_2_ and temperature-controlled chamber. For mitochondrial transport, axons were imaged in widefield with a 60× oil objective for 5 min at 10 s intervals. Kymographs were generated in FIJI and trajectories were analysed with KymoButler (Jakobs et al., 2019).

Post-processing in FIJI included uniform background subtraction (25-pixel radius), smoothing, and contrast enhancement. Quantification of morphology, translation, and apoptosis was done using CellProfiler (v4.2.8) with custom pipelines. IdentifyPrimaryObjects module was used with an adaptive two-class Otsu threshold (smoothing scale 1.34). Object diameter settings: nuclei (DAPI), 15–60 pixels; mitochondria (TOMM20 or MitoBrilliant), 2–20 pixels; apoptotic nuclei (TUNEL), 5–40 pixels; translating mitochondria (HPG), 0.5–20 pixels. Clumped nuclei/mitochondria were classified by intensity and separated by shape. Mitochondria were further filtered (FilterObjects) to retain those with a form factor between 0.3 - 1.5 and area ≤ 300 pixels. To threshold dendritic areas, MAP2-positive neurites were enhanced using the module EnhanceOrSuppressFeatures (tubeness method, smoothing scale = 2), followed by Minimum-Cross-Entropy thresholding and size selection (20-40 pixels). Object relations and masks were derived with RelateObjects/MaskObjects and measurements used MeasureObjectSizeShape.

High-throughput imaging was performed on the Opera Phenix system. Images of the neurons were obtained at 20x and 40x magnification. Images were processed on Columbus (v2.9.1) with custom pipelines. The command Find Nuclei (Method C) was used to identify nuclei, stained by Hoescht. The command Find Spots (Method B) was used to identify mitochondria (MitoBrilliant), and Calculate Morphology Properties was used to identify the area of each individual mitochondria. Mitochondria within the cell body, which could not be individually resolved, were removed from analysis using Select Population – where identified spots over 150px^2^ were removed from consideration.

#### Transmission electron microscopy (TEM)

Neurons were fixed in 4% PFA/2.5% glutaraldehyde in 0.1 M sodium cacodylate (20 min, RT), post-fixed in 1% osmium tetroxide/1.5% potassium ferricyanide (1 h, RT), washed, and stained in 1% aqueous uranyl acetate overnight. Samples were dehydrated (50–70–95–100% ethanol; 10 min each; 100% ×3).

Embedding was done using TAAB Epoxy Resin 812 (formulations per manufacturer), with graded infiltration (25% 3 h, 50% overnight, 75% 3 h, then 100% overnight). Resin-filled capsules were polymerized overnight. Ultrathin sections (∼70 nm) were cut with a diamond knife and collected on copper grids. Sections were stained with UA Zero (20% ethanol, 1 min) and lead citrate (30 s), rinsed, and imaged on a Hitachi TEM with Xarosa camera at 5,000x, 10,000x and 20,000x.

#### Luminescence-based metabolic assays

All assays used Promega kits and were read on a GloMax microplate reader (integration 0.3 s), with signals normalised to cell number.

**Lactate (Lactate-Glo, J5021)**: conditioned media were collected 2 h after a fresh medium change and mixed 1:1 with Lactate Detection Reagent (1 h, RT). Medium incubated without cells served as background.

**Glucose uptake (Glucose Uptake-Glo, J1342):** live neurons were incubated with 1 mM 2-DG (20 min, RT), then reactions were stopped with Stop and Neutralization buffers (kit), which also lysed the cells. Lysates were then incubated with 2DG6P Detection Reagent (1 h, RT). A no-2DG control was included.

**ATP (CellTiter-Glo, G7571):** this assay was multiplexed with the glucose uptake assay according to the manufacturer’s instructions. Following the glucose assay stop step, 10 µL lysate was mixed with 200 µL CellTiter-Glo reagent (10 min, RT) and luminescence recorded.

#### Bulk RNA-sequencing and analysis

Neuronal cultures were gently dissociated with papain (6 min, 37°C). Total RNA (mirVana kit, Thermo Fisher) was DNase-treated (Bio-Rad). Libraries were prepared by the Wellcome-MRC Cambridge Stem Cell Institute (poly(A) selection, Takara SMARTer Stranded Total RNA-Seq) and sequenced on an Illumina NovaSeq S1 (PE50), ∼26 M reads/sample (n=3 biological replicates/condition).

Sequencing quality was assessed with FastQC, and adapter and low-quality reads were removed using Trim Galore! (v0.6.5). Reads were aligned to GRCh38 with STAR (v2.7.3a). Gene-level counts were obtained with featureCounts, and differential expression analysis was performed with DESeq2 with Wald tests and Benjamini–Hochberg correction (FDR<0.05). Both a simple model (∼condition) and an interaction model (∼condition * cell type) were used. Variance-stabilized counts (VST) were used for visualisation. Functional enrichment analysis was conducted using clusterProfiler for gene set enrichment analysis (GSEA), and additional pathway analyses were performed using Enrichr-KG.

#### Statistical Analysis

Statistical analyses were performed using GraphPad Prism v10 and R Studio v4.3.1. Data are presented as mean ± standard error of the mean (SEM) unless stated otherwise. Biological replicates correspond to independent differentiation rounds, and technical replicates were averaged within each differentiation. Data from different sgRNA lines were pooled and treated as independent biological replicates unless specified otherwise in the figure legends.

Parametric tests were used after confirming that data distributions did not significantly deviate from normality, as assessed by Shapiro-Wilk tests and visual inspection of Q-Q plots. Homogeneity of variance was evaluated using F-tests, with Welch’s correction applied where variances were unequal. Comparisons between two independent groups were performed using unpaired two-tailed Student’s t-tests, and comparisons involving more than two groups were analysed using one-way ANOVA followed by Tukey’s multiple-comparison test. Statistical significance was set at p < 0.05.

For bulk RNA-seq, DESeq2 was used with FDR<0.05 and |log_2_ fold change| ≥ 0.5. Outliers were not excluded given small sample sizes and intrinsic biological variability and all data points were retained unless clear technical errors were identified. Imaging/molecular/luminescence plots were generated in Prism, transcriptomics plots used ggplot2 and related packages in R. Significance is annotated as \**p<0.05, **p<0.01, ***p<0.001, ****p<0.0001*.

### ETHICAL ISSUES

We used human primary fibroblasts to generate induced pluripotent stem cells (iPSCs) and then differentiate them to 2D neuronal cells. Participants have been consented for using their cells for this research as part of the clinical research study “Genotype and phenotype in inherited neurodegenerative diseases” (REC ID: 13/YH/0310).

## KEY RESOURCE TABLE

| Reagent or resource | Source | Identifier |
| --- | --- | --- |
| **Antibodies** |  |  |
| ISLET-1 | Abcam | Cat# ab109517 |
| BIII-tubulin | Abcam | Cat# ab78078 |
| BIII-tubulin | Aves Labs | Cat# TUJ-0020 |
| MAP2 | Abcam | Cat# ab5392 |
| MAP2 | Thermo Fisher Scientific | Cat# PA1-10005 |
| SMI132 | BioLegend | Cat# 837904 |
| CTIP2 | Abcam | Cat# ab18465 |
| ChAT | Invitrogen | Cat# CL3173 |
| ChAT | Sigma | Cat# AB143 |
| vGlut1 | Synaptic Systems | Cat# 135 303 |
| TOMM20 | Abcam | Cat# ab186735 |
| TOMM20 | Proteintech | Cat# 11802-1-AP |
| MT-CO1 | Abcam | Cat# ab14705 |
| Vinculin | Santa Cruz | Cat# sc-73614 |
| TFAM | Cell Signalling | Cat# D5C8 |
| dsDNA | Abcam | Cat# ab27156 |
| Total OXPHOS Human WB Cocktail Kit | Abcam | Cat# ab110411 |
| Donkey Anti Goat IgG (H+L) Highly Crossed Absorbed Secondary Antibody, Alexa Fluor Plus 647 | Thermo Fisher Scientific | Cat# A32849TR |
| Donkey Anti Mouse IgG (H+L) Highly Crossed Absorbed Secondary Antibody, Alexa Fluor 568 | Thermo Fisher Scientific | Cat# A10037 |
| Donkey Anti Rabbit IgG (H+L) Highly Crossed Absorbed Secondary Antibody, Alexa Fluor 488 | Thermo Fisher Scientific | Cat# 21206 |
| Donkey Anti Rabbit IgG (H+L) Highly Crossed Absorbed Secondary Antibody, Alexa Fluor 568 | Thermo Fisher Scientific | Cat# SA5-10042 |
| Donkey Anti Rat IgG (H+L) Alexa Fluor 647 preabsorbed | Abcam | Cat# ab150155 |
| Goat Anti Chicken IgY (H+L) Cross-Absorbed Secondary Antibody Alexa Fluor 647 | Thermo Fisher Scientific | Cat# A48260 |
| Donkey Anti Mouse IgG (H+L) Highly Crossed Absorbed Secondary Antibody, Alexa Fluor 647 | Thermo Fisher Scientific | Cat# A-31571 |
| Goat anti-Chicken IgY (H+L) Secondary Antibody, Alexa FluorTM 568 | Thermo Fisher Scientific | Cat# A-11041 |
| Goat Anti Mouse IgG (H+L) Secondary Antibody, HRP | Thermo Fisher Scientific | Cat# G-21040 |
| Goat Anti Rabbit IgG (H+L) Secondary Antibody, HRP | Thermo Fisher Scientific | Cat# 31462 |
| MitoBrilliant 646 | Tocris | Cat# 7700 |
| Janelia Fluor HaloTag Ligand, JFX 650, lyophilized | Promega | Cat# HT1070 |
| **Chemicals, peptides and Recombinant Proteins** |  |  |
| DAPI Nuclear Counterstain | Thermo Fisher Scientific | Cat# 62248 |
| Accutase | Sigma-Aldrich | Cat# A1110501 |
| Antibiotic-Antimycotic (100X) | Thermo Fisher Scientific | Cat# A86744-25MG |
| Bovine Serum Albumin | Thermo Fisher Scientific | Cat# A9418-100G |
| BrainPhys Neuronal Medium | STEMCELL Technologies | Cat# 5790 |
| Complete, Mini, EDTA free Protease Inhibitor Cocktail | Sigma-Aldrich | Cat# 11836170001 |
| Compound E | STEMCELL Technologies | Cat# 73952 |
| Cytiva's Amersham ECL Select Western Blotting Detection Reagent | Cytiva | Cat# RPN2235 |
| DMSO | Sigma-Aldrich | Cat# D2650 |
| Dpbs, no calcium, no magnesium | Thermo Fisher Scientific | Cat# 14-190-144 |
| Doxycycline hyclate | Sigma-Aldrich | Cat# D9891-1G |
| Essential 8 Medium | Thermo Fisher Scientific | Cat# A1517001 |
| Geltrex | Thermo Fisher Scientific | Cat# A1413302 |
| Gibco DMEM/F12, GlutaMAX | Thermo Fisher Scientific | Cat# 10-565-018 |
| DMEM/F12 | Thermo Fisher Scientific | Cat# 11320033 |
| Neurobasal Medium | Thermo Fisher Scientific | Cat# 21103049 |
| GlutaMAX | Thermo Fisher Scientific | Cat# 35050061 |
| NEAA | Thermo Fisher Scientific | Cat# 11140050 |
| B27 | Thermo Fisher Scientific | Cat# 17504044 |
| Homopropargylglycine (HPG) | Thermo Fisher Scientific | Cat# C10186 |
| Laminin | Corning | Cat# 354239 |
| Lipofectamine Stem Transfection Reagent | Thermo Fisher Scientific | Cat# STEM00003 |
| Molecular Probes ProLong Gold Antifade Mountant | Thermo Fisher Scientific | Cat# P36934 |
| N-2 Supplement (100X) | Thermo Fisher Scientific | Cat# 17502048 |
| NeuroCult SM1 Neuronal Supplement | STEMCELL Technologies | Cat# 5711 |
| Opti-MEM Reduced Serum Medium | Thermo Fisher Scientific | Cat# 31985062 |
| Pierce Methanol-free Formaldehyde Ampules | Thermo Fisher Scientific | Cat# 28908 |
| Pierce Rapid Gold BCA Protein Assay Kit | Thermo Fisher Scientific | Cat# A53225 |
| Purmorphamine | Sigma-Aldrich | Cat# 540220-5MG |
| Puromycin dihydrochloride | Sigma-Aldrich | Cat# P8833-25MG |
| G418 | Sigma-Aldrich | Cat# G8168 |
| Polybrene | Sigma-Aldrich | Cat# TR-1003-G |
| Recombinant Human BDNF | PeproTech | Cat# 450-02 |
| Recombinant Human GDNF | Bio-Techne | Cat# 212-GD-010 |
| Recombinant Human NT-3 | PeproTech | Cat# 450-03-10ugmito |
| Human recombinant BDNF | STEMCELL Technologies | Cat# 78005 |
| Human recombinant GDNF | STEMCELL Technologies | Cat# 78058 |
| Human recombinant CNTF | STEMCELL Technologies | Cat# 78010 |
| Cytarabine (Ara-C) | Sigma-Aldrich | Cat# C6645-25MG |
| ReLeSR | STEMCELL Technologies | Cat# 100-0483 |
| Smoothened Agonist SAG-CAS | Sigma-Aldrich | Cat# 566660-1MG |
| Smoothened Agonist (SAG) | STEMCELL Technologies | Cat# 73412 |
| SB431542 | STEMCELL Technologies | Cat# 72232 |
| Retinoic Acid | Sigma-Aldrich | Cat# R2625-50MG |
| LDN193189 | STEMCELL Technologies | Cat# 72147 |
| Poly-D-Lysine | Thermo Fisher Scientific | Cat# A3890401 |
| Triton X-100 | Sigma-Aldrich | Cat# T8787 |
| Y-27632 dihydrochloride | Tocris | Cat# 1254 |
| 2'-Deoxy-5-ethynyluridine | Biosynth | Cat# NE08701 |
| **DNA Constructs** |  |  |
| pKLV-U6gRNA(BbsI)-PGKpuro2ABFP | Sanjana et al. | Addgene #52963 |
| CMV-mito-R-GECO1, | Wu et al. | Addgene #46021 |
| pAAV.CAG.(mito).iATPSnFR2.A95A.A119L.HaloTag | Marvin et al. | Addgene #209722 |
| Packaging plasmid p8.91 | Susac et al. | Addgene #187441 |
| Envelope plasmid pMD.G | Susac et al. | Addgene #187440 |
| pCXLE-hOct3/4 shp53-F | Okita et al. | Addgene #27077 |
| pCXLE-hSK | Okita et al. | Addgene #27078 |
| pCXLE-hUL | Okita et al. | Addgene #27080 |
| pCXLE-eGFP | Okita et al. | Addgene #27082 |
| FUdeltaGW-rtTA | Maherali et al. | Addgene #19780 |
| pTet-O-Ngn2-puro | Zhang et al. | Addgene #52047 |
| **Experimental models: cell lines** |  |  |
| Human iPSCs expressing dCas9-KRAB and TetO-NGN2 | Gift from Dr. Evan Reid, University of Cambridge | N/A |
| PGPC-3, whole genome sequenced control line | Gift from Dr. James Ellis (Hildebrandt et al.) | N/A |
| **Software** |  |  |
| FIJI |  | RRID:SCR_002285 |
| CellProfiler 4.2.8 | https://cellprofiler.org | RRID:SCR_007358 |
| R Studio |  | RRID:SCR_000432 |
| GraphPad Prism 10 |  | RRID:SCR_002798 |
| Biorender | https://biorender.com | RRID:SCR_018361 |
| Columbus |  | RRID:SCR_025391 |
| **Kits/ Other** |  |  |
| NuPAGE Novex 4-12% Bis-Tris Protein Gels | Thermo Fisher Scientific | Cat# NP0335BOX |
| NuPAGE MES SDS Running Buffer | Thermo Fisher Scientific | Cat# NP0002 |
| iBlot 2 Transfer Stacks, PVDF, mini | Thermo Fisher Scientific | Cat# IB24002 |
| ddPCR Supermix for Probes (no dUTP) | Bio-Rad | Cat# 1863023 |
| IscripT gDNA cDNA Syntehsis Kit | Bio-Rad | Cat# 1725035 |
| Molecular Probes Click iT Cell Reaction Buffer Kit | Thermo Fisher Scientific | Cat# C10269 |
| Glucose Uptake-Glo Assay | Promega | Cat# J1341 |
| CellTiter-Glo Luminescent Cell | Promega | Cat# G7570 |
| Lactate-Glo Assay | Promega | Cat# J5021 |
| One Shot Stbl3 Chemically Competent | Thermo Fisher Scientific | Cat# C737303 |
| pPACKH1 HIV Lentivector Packaging Kit | System Biosciences | Cat# LV500A-1 |
| PEG-it Virus Precipitation Solution | System Biosciences | Cat# LV810A-1 |

