## Supplementary figures and images for "Cell-type-specific adaptations to mitochondrial stress underly the neurological presentations of *MTRFR* mutations"

### Supplementary Figures

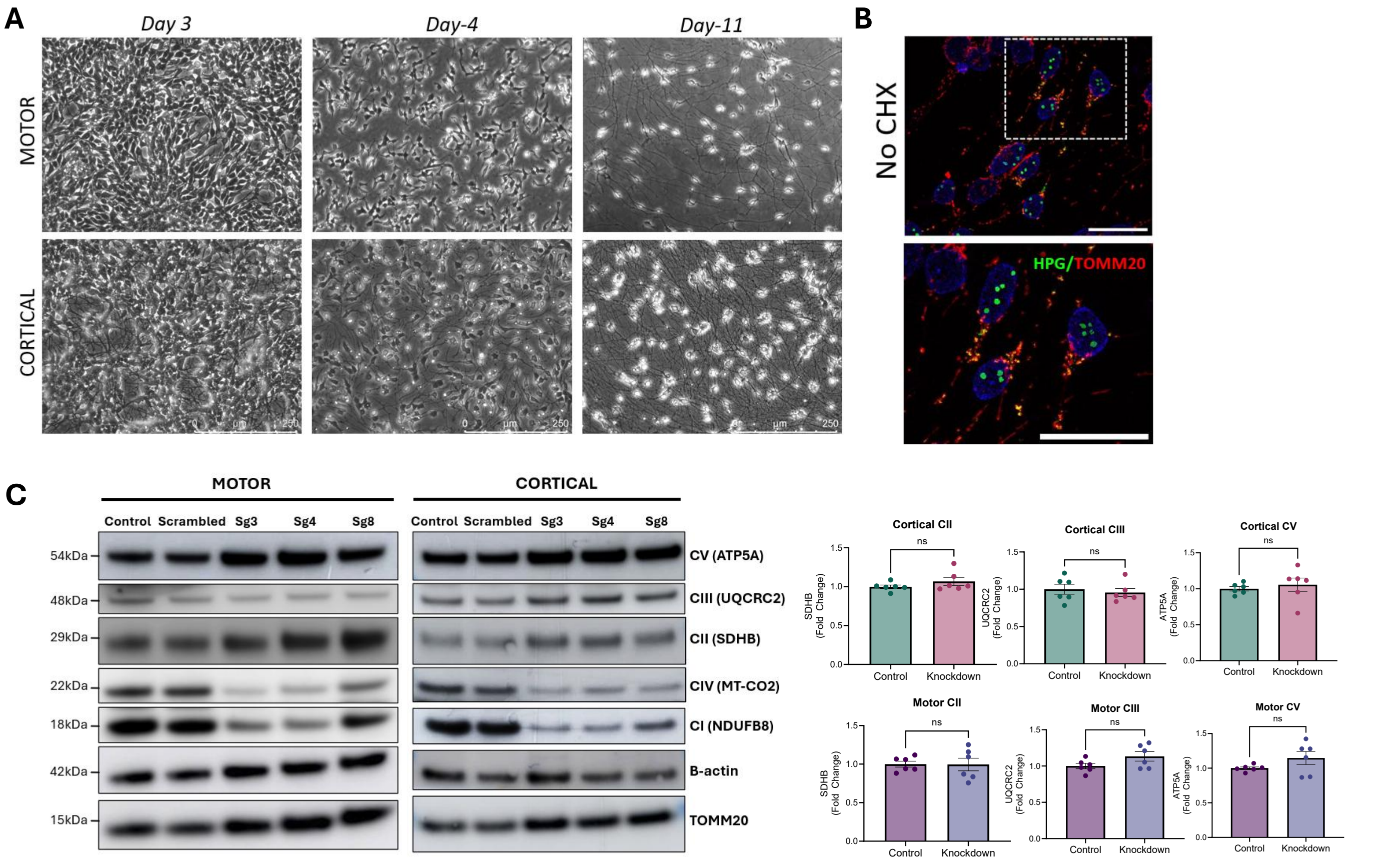

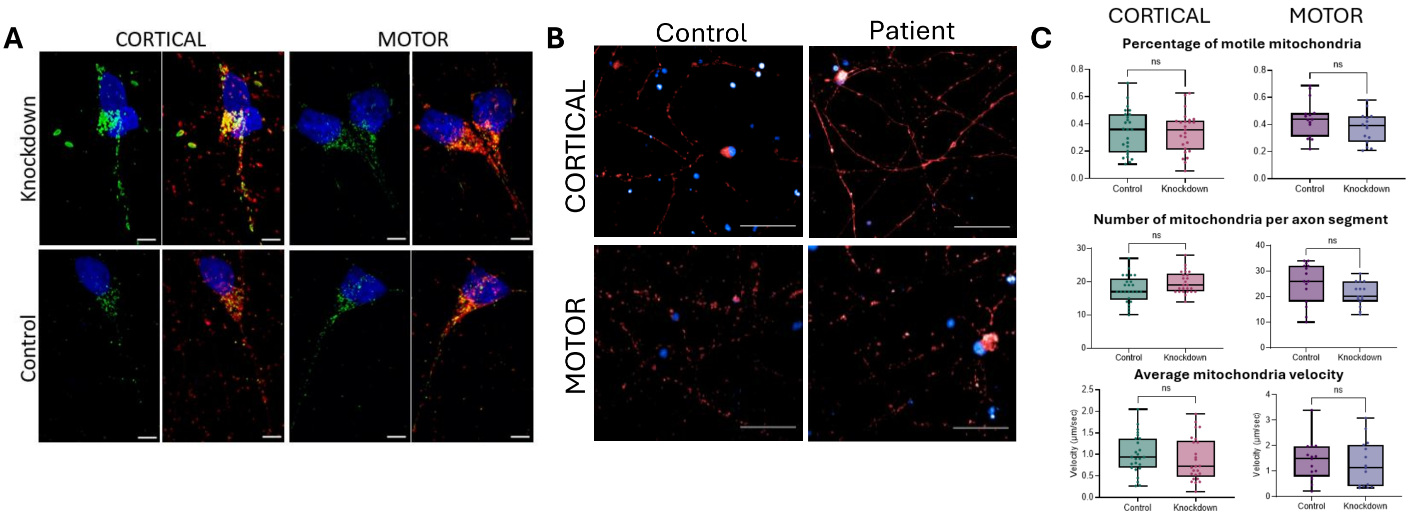

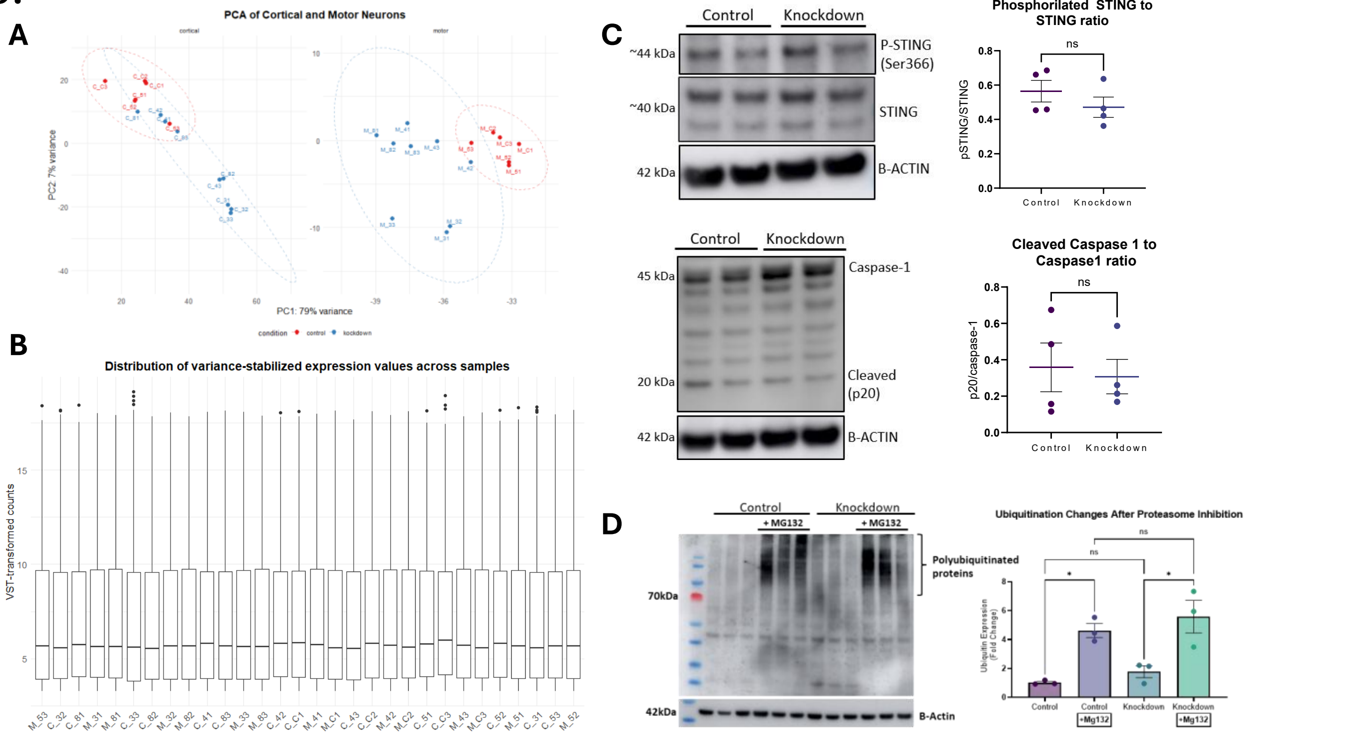
