## Supplementary material for "Cell-type-specific adaptations to mitochondrial stress underly the neurological presentations of *MTRFR* mutations": Table S1

| Single guide name | Target sequence (5’-3’) | Oligo sequence; sense (5’-3’) | Oligo sequence; antisense (5’-3’) |
|---|---|---|---|
| **KD1** | TATCCAAATCACCTCAGCGG | CACCGTATCCAAATCACCTCAGCGG | AAACCCGCTGAGGTGATTTGGATAC |
| **KD2** | TCGCAAGTCGGGAAAAACCC | CACCGTCGCAAGTCGGGAAAAACCC | AAACGGGTTTTTCCCGACTTGCGAC |
| **KD3** | CGATTGCGAATCCTCCGCTG | CACCGCGATTGCGAATCCTCCGCTG | AAACCAGCGGAGGATTCGCAATCGC |
| **Scramble** | GGGACGCGAAAGAAACCAGT | CACCGGGACGCGAAAGAAACCAGT | AAACACTGGTTTCTTTCGCGTCCC |
