## Supplementary material for "Cell-type-specific adaptations to mitochondrial stress underly the neurological presentations of *MTRFR* mutations": Table S2

| Target | Forward (5’ to 3’) | Reverse (5’ to 3’) |
|---|---|---|
| *MTRFR*-exon1 | GAGAGAGGACTGCGAGCC | CGTAAACCAGGTCCCGAGAA |
| *MTRFR*-exon2 | CCTACACCACTGACCCGAAT | TAGTCCTTCTTGCCTGCCAT |
| *β-ACTIN* | CACCATTGGCAATGAGCGGTTC | AGGTCTTTGCGGATGTCCACGT |
| *HSPA1B* | ACCTTCGACGTGTCCATCCTGA | TCCTCCACGAAGTGGTTCACCA |
| *HSP90AA1* | TCTGCCTCTGGTGATGAGATGG | CGTTCCACAAAGGCTGAGTTAGC |
| *HSF1* | TGAAAAGTGCCTCAGCGTAGCC | TGCTCAGCATGGTCTGCAGGTT |
| *DNAJB1* | AGTTCAAGGAGATCGCTGAGGC | GCTGAAAGAGGTACCATTGGCAC |
| *MT-ND1* | GGGTTCATAGTAGAAGAGCGATGG | ACGCCATAAAACTCTTCACCAAAG |
| *MT-ND4* | AGTGCGATGAGTAGGGGAAGG | ACCTTGGCTATCATCACCCGAT |
| *B2M* | CGCAATCTCCAGTGACAGAA | GCAGAATAGGCTGCTGCTGCTGTTCC |
| *RNaseP* | AGATTTGGACCTGCGAGCG | GAGCGGCTGTCTCCACAAGT |
